# Ultra-deep sequencing reveals intra-host diversity and co-infection-driven evolution of SARS-CoV-2

**DOI:** 10.64898/2026.01.15.699801

**Authors:** Carol Moraga, Francisco Kirhman, Barbara Bernal, Sofia Poblete, Pastor Jullian, Adrian Gonzalez, Mauricio Latorre, Alex Di Genova

## Abstract

As COVID-19 enters an endemic phase, SARS-CoV-2 continues to diversify under ongoing immune pressure, with Omicron sublineages and episodic emergent variants sustaining reinfections worldwide. Intra-host evolution represents the earliest stage of this diversification, yet remains undercharacterized, particularly in regions with limited genomic surveillance. Here, we conducted high-throughput sequencing on 96 nasopharyngeal swab samples from Chilean individuals (2020–2022), achieving an average per-base genome coverage of ∼60,000x across the viral genome. This ultra-deep sequencing coverage enabled the identification of intra-host single-nucleotide variants (iSNVs) and co-infection events with high sensitivity and accuracy. Co-infections, especially with Omicron, significantly increased iSNV frequency and recombination, driving viral diversity. Evolutionary analysis based on the non-synonymous to synonymous ratio (dN/dS) shows that Omicron is under extensive purifying selection (global dN/dS ∼ 0.55). However, Omicron co-infection cases exhibited higher dN/dS ratios (∼0.58), suggesting a lower level of purifying selection and increased genetic diversity. Notably, the Spike gene showed dN/dS ratios indicative of positive selection (dN/dS > 1), which are more pronounced in co-infection cases than in Omicron alone. This suggests that co-infections are providing the substrate for the emergence of new variants with enhanced transmissibility and immune evasion capabilities. Together, these findings demonstrate that ultra-deep sequencing is crucial for mapping the evolutionary forces driving SARS-CoV-2 intra-host adaptation and the emergence of new variants.

## Introduction

The SARS-CoV-2 pandemic has resulted in more than 778 million confirmed cases and more than 7 million deaths worldwide[1]. Although global transmission and disease severity have decreased and COVID-19 has transitioned toward endemicity[2], the virus continues to evolve[3], generating recurrent waves[4,5] driven by waning immunity and the emergence of immune-evasive variants[6,7]. Between 2021 and 2023, the Omicron variant and its numerous sublineages significantly reshaped the global epidemiological landscape [4–9]. Omicron rapidly displaced earlier variants due to its high transmissibility, extensive spike protein mutations, and enhanced capacity for reinfection, even in individuals with prior infection or vaccination [2,4–8].

Chile implemented a comprehensive strategy that combined large-scale viral detection, efficient booster-dose administration, and targeted quarantine strategies, becoming one of the world’s leading nations in the rapid deployment of SARS-CoV-2 vaccination programs and ultimately immunizing more than 90% of the target population within a remarkably short period [10–13]. However, structural inequities in healthcare access, particularly in lower-income municipalities, were reflected in elevated infection fatality rates throughout the pandemic [14]. Within this context, the period between 2020 and 2022 in Chile, spanning the transition from early lineages to Omicron dominance [11, 12], provides a valuable retrospective window for studying how SARS-CoV-2 evolved during pivotal phases of the pandemic.

Genome sequencing was central to global SARS-CoV-2 surveillance efforts[15], supported by large-scale data sharing platforms such as GISAID, which surpassed 20 million sequences by 2025[16], and by complementary genotyping approaches that provide rapid detection of variant-defining mutations[17,18]. While most surveillance has focused on consensus-level mutations across individuals[19–21], intra-host viral evolution, the earliest stage of SARS-CoV-2 diversification, remains comparatively understudied[19,20,22,23], particularly in underrepresented regions such as Chile. SARS-CoV-2, like other RNA viruses, accumulates mutations within infected individuals due to replication errors, RNA damage, and host-mediated editing by enzymes such as APOBEC and ADAR[24]. These processes generate intra-host single-nucleotide variants (iSNVs), typically defined within a variant allele frequency range of 5–95%[20]. Detecting iSNVs reliably requires ultra-deep sequencing, often exceeding 1000× and ideally reaching tens of thousands of coverage, to distinguish true low-frequency variants from sequencing artifacts[20,23]. Intra-host variation offers important insight into the microevolutionary processes that generate diversity, shape transmission fitness, and may give rise to emerging variants[19–22,25,26]. Co-infections, where two or more SARS-CoV-2 variants simultaneously infect an individual, represent an additional source of elevated intra-host diversity and may create opportunities for recombination and accelerated viral evolution[3,26]. Such events are expected to be more frequent during periods when divergent lineages co-circulate, such as the Chilean transitions from Gamma and Lambda to Omicron in 2020–2022. Yet, systematic characterization of co-infections and their evolutionary consequences remain limited[26].

Here, we present the first ultra-deep SARS-CoV-2 whole-genome sequencing analysis of intra-host evolution in Chile, using the MGI ATOPlex platform[27] and DNBSeq-G400[28] sequencer to generate exceptionally high coverage (∼60,000× on average) across 96 samples collected between 2020 and 2022. This approach enabled sensitive detection of low-frequency iSNVs and robust identification of co-infections. By integrating consensus-level and intra-host analyses, we characterized mutational spectra, quantified iSNV burdens, identified co-infection events, reconstructed lineage relationships, and estimated global and gene-level dN/dS ratios across major SARS-CoV-2 variants circulating during this period.

Our study provides novel insight into the within-host evolutionary dynamics of SARS-CoV-2 across successive epidemic waves in Chile, revealing the influence of lineage background and co-infection on intra-host diversity, identifying gene-specific episodes of positive selection, and demonstrating how ultra-deep sequencing can uncover early evolutionary processes that may contribute to the emergence of future variants.

## Results

### SARS-CoV-2 genome sequencing of Chilean samples from 2020-2022

We sequenced 96 COVID-19 samples from Chile (2020–2022) at an average depth of ∼60,000x, enabling high-confidence detection of intra-host variants. Whole-genome virus sequencing was performed on a DNBSeq-G400[28] sequencer using the ATOPlex method[29] from nasopharyngeal (NP) swabs of confirmed COVID-19 individuals from two Chilean regions (Figure S1). The multiplex PCR of ATOPlex utilizes short amplicons (106-199 bp) with high specificity and sensitivity, enabling the generation of complete SARS-CoV-2 virus genomes from samples with PCR Ct values of up to 35 [27,29].

Sequencing generated a total of 2,090 million paired-end 100 bp reads for the 96 COVID-19 samples. The sequencing data exhibited high quality across several metrics (Figure 1). All sequencing data were analyzed using the CECRET[30] Nextflow[31] pipeline, which includes quality control, mapping, variant calling, and clade classification (see Materials and Methods). The PCR cycle threshold (Ct) values varied across samples, with a median of 18 and a range of 10–37 (SD = 6.21), reflecting differences in viral RNA abundance in the starting material (Figure 1). As viral load is inversely proportional to Ct[32], lower values indicate higher concentrations of viral RNA and, consequently, greater infectiousness. The total reads per sample had a median of 21.8 million, ranging from 3.62 million to 55.3 million (SD = 11.1 million). The average read quality (Q-score) was 35.3, with minimal variation (SD = 0.160), indicating consistently high-quality reads. The Mapping of the reads to the Wuhan-Hu-1 (RefSeq ID NC_045512) reference was high, with 99.8% of reads mapped (Figure 1). The read error rate was low, with a median of 0.00672%. These metrics indicate that our in-house COVID-19 sequencing, utilizing ATOPLEX and the DNBSeq-G400 sequencer, generated high-quality reads and achieved a high mapping rate for the 96 samples, despite some variability in cycle threshold (Ct) values (Figure 1).

**Figure 1:**
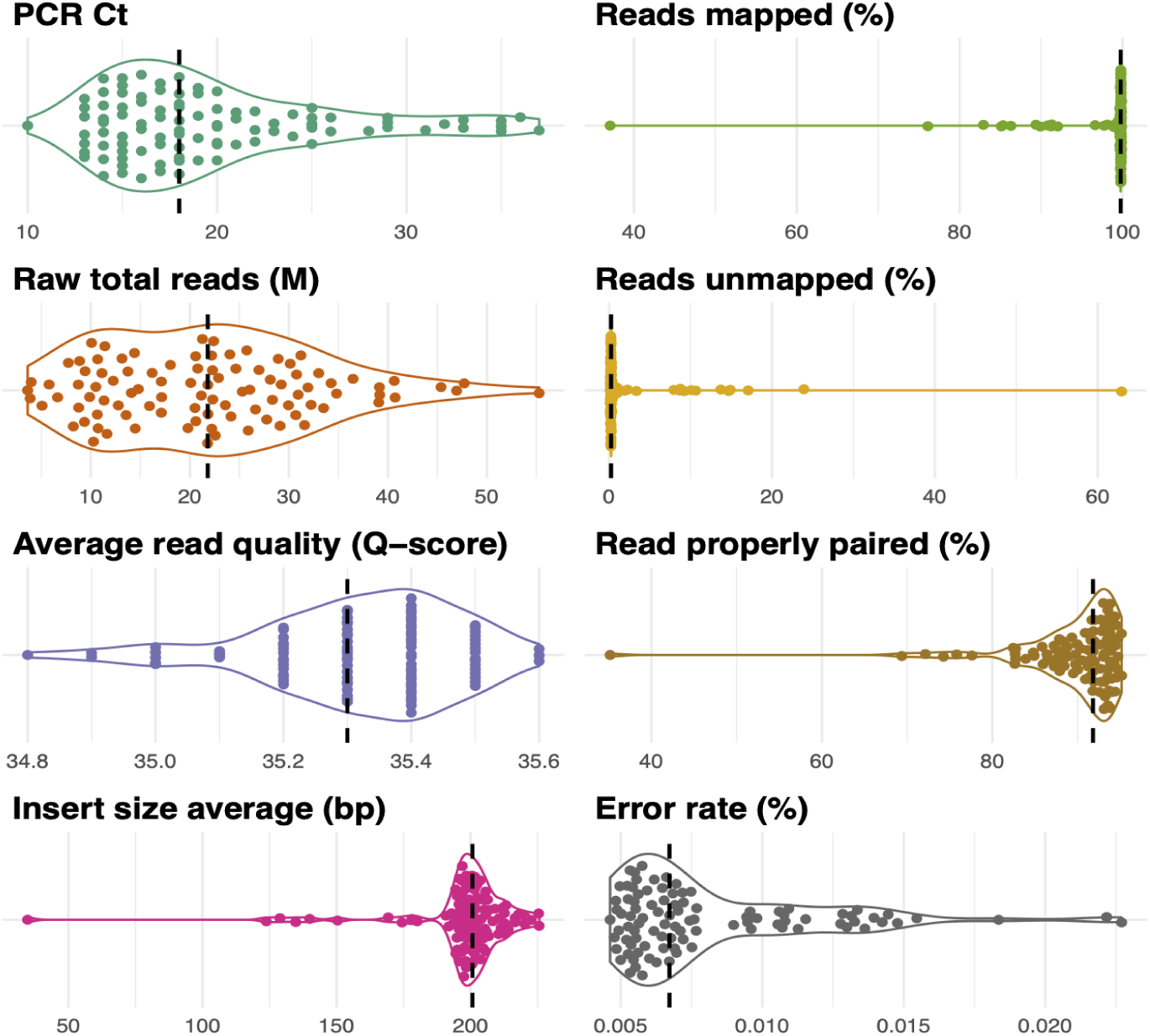
Quality Metrics for 96 Sequenced COVID-19 Samples. Panels include PCR cycle threshold (Ct) values, total reads sequenced in millions, average read quality scores (Q-scores), and insert size averages in base pairs. Additionally, the percentages of reads mapped, reads unmapped, and reads properly paired are shown, along with the error rates. Each dot represents a sequenced sample. Each panel displays the distribution (violin plot), median (black dotted line), and variability of these metrics across the dataset, illustrating the overall high quality of the sequencing data.

Our data significantly contribute to Chilean SARS-CoV-2 genomic data, which showed fewer than 1,000 COVID-19 genomes sequenced monthly during 2020–2021 (Figure S1C). Chilean SARS-CoV-2 genome sequences deposited on GISAID (Figure S1C) demonstrate a variation in prevalence over the study period, consistent with global trends (Figure S1C). In 2020, unidentified or pre-existing variants categorized as “Other” were most abundant. The Alpha variant, first detected globally in late 2020, emerged in the Chilean samples in 2021, and it was then subsequently overtaken by the Delta and finally by the Omicron variant, it surfaced in 2021 and rapidly rose to become the dominant variant by 2022, aligning with its swift global dissemination (Figure S1C). The sequenced samples, set against the backdrop of Chilean genomes submitted to GISAID from 2020 to 2023 (45,466 genomes as of 22 September 2023), highlight the contribution of our data to Chilean efforts to build SARS-CoV-2 genomic datasets. This period encompasses significant Variants of Concern (VOCs), including Lambda, Delta, and Omicron (Figure S1C).

### Phylogenetic analysis and sequencing depth

Phylogenetic analysis revealed distinct clustering of Omicron (BA.2.X), Gamma, Alpha, and Lambda variants, with Omicron showing the highest divergence (Figure 2). Out of the 96 samples, 40 belonged to the Omicron (BA.2.X) variant, which harbored the highest average number of SNPs per sample (mean = 65). The Gamma variant, represented by 20 samples, followed with an average of 45 SNPs per sample, indicating substantial but less extensive diversification. In contrast, the Alpha variant, with 15 samples, and the Lambda variant, with 10 samples, exhibited lower genetic variability, with averages of 25 and 22 SNPs per sample, respectively (Figure 2). Epsilon, which included 5 samples, showed minimal divergence with the lowest SNP count across samples. These results are consistent with global trends in SARS-CoV-2 evolution[3], where Omicron and its sub-lineages have shown the highest levels of genetic diversity and rapid mutation, contributing to their widespread prevalence and ongoing evolution.

**Figure 2.**
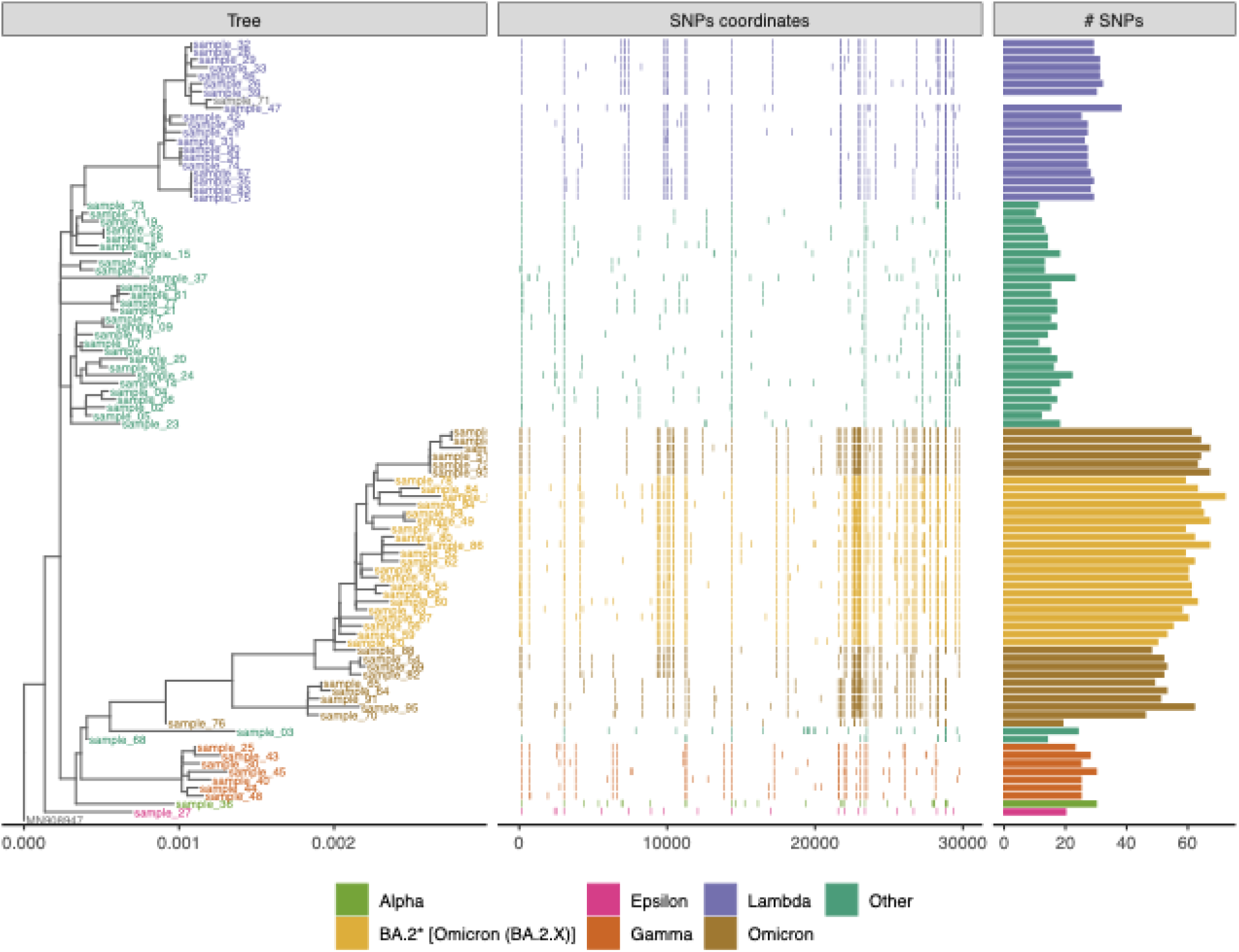
Phylogeny, genome-wide SNP distribution, and mutational burden of Chilean SARS-CoV-2 genomes. The maximum-likelihood tree (left) separates Alpha, Gamma, Lambda, Omicron (BA.2*), Epsilon, and Other lineages. The central panel shows SNP positions along the SARS-CoV-2 genome for each sample, ordered by the phylogeny. The right panel shows total SNP counts, with Omicron exhibiting the greatest divergence from earlier variants.

A distinctive feature of our sequencing effort is the ultra-deep genome coverage achieved, enabling, for the first time, a comprehensive characterization of intra-host single-nucleotide variation in SARS-CoV-2 samples from Chilean individuals. Such ultra-high coverage enables the reliable detection of low-frequency alleles, providing a foundation for investigating early viral diversification and adaptation within hosts[3,21,22,25]. Within-host mutational patterns differ from population-level SNVs because they capture early, transient stages of viral replication and editing. Across all samples, coverage ranged from 2,980× to 442,413× per position, with an average depth exceeding 47,000× per variant (Supplementary Fig. 2), ensuring the coverage and completeness required for downstream intra-host variant analysis[20,23].

### Determinants of intra-host SARS-CoV-2 genetic diversity reveal a predominant effect of co-infections

To ensure reliable identification of iSNVs, we applied stringent quality filters to the sequencing data. Variants were required to have an alternative allele depth of at least 100 reads, a quality score above 30, and a frequency between 5% and 95% to exclude potential sequencing artifacts and fixed variants[20]. Positions overlapping known consensus SNVs in the same sample were excluded to avoid redundancy. Only high-quality samples with cycle threshold (Ct) values ≤ 28[33] were retained for downstream analysis, and cases showing >3% representation of an additional lineage were classified separately as potential co-infections.

Analysis of 96 SARS-CoV-2 genomes revealed distinct temporal patterns in SNVs and iSNVs between 2020 and 2022 (Supplementary Figure 3). SNV counts increased significantly over time (R² = 0.83, p < 0.001), reflecting the accumulation of fixed mutations across epidemic waves, particularly in 2022 during Omicron dominance. In contrast, iSNV frequencies remained relatively stable (R² = 0.08), indicating that within-host mutational diversity did not markedly change over the study period. We identified 19 samples (≈20%) exhibiting evidence of co-infection, each containing between 2 and 5 distinct SARS-CoV-2 lineages (Supplementary Table 1). The most frequent combinations involved Omicron sublineages (BA.2* and BA.5*) co-occurring with Other, Gamma, or Lambda variants. Notably, the most complex cases contained up to five coexisting variants within a single sample, indicating extensive viral mixing during the Chilean epidemic transition from Gamma/Lambda to Omicron dominance.

We modeled the number of iSNVs using a linear model with age group, sex, mutation load (iVar), variant, PCR Ct, and average genome coverage (log) as predictors (Figure 3). The model explained ∼47% of the variance (R² = 0.472, adjusted R² = 0.383; F(12, 71) = 5.30, p = 2.70×10⁻⁶). No single coefficient reached significance after adjustment, although the effect of male sex trended lower iSNV counts (β = −3.25 for male vs reference; p = 0.078, Figure 3B). This trend can be attributed to differences in immune responses during the course of SARS-CoV-2 infection in man and female patients [34]. Neither Ct values (β = 0.44, SE = 0.28, p = 0.12) nor average genome coverage (β = 0.58, SE = 3.12, p = 0.85) were associated with iSNV counts, suggesting that technical variation and viral load within the studied range did not drive the observed diversity differences across samples(Figure 3A). Co-infected cases showed a positive but imprecise increase in iSNVs (β = 9.75, p = 0.27), consistent with a possible elevation in diversity when multiple viral lineages co-occur[3], albeit with considerable uncertainty (Figure 3A).

**Fig. 3:**
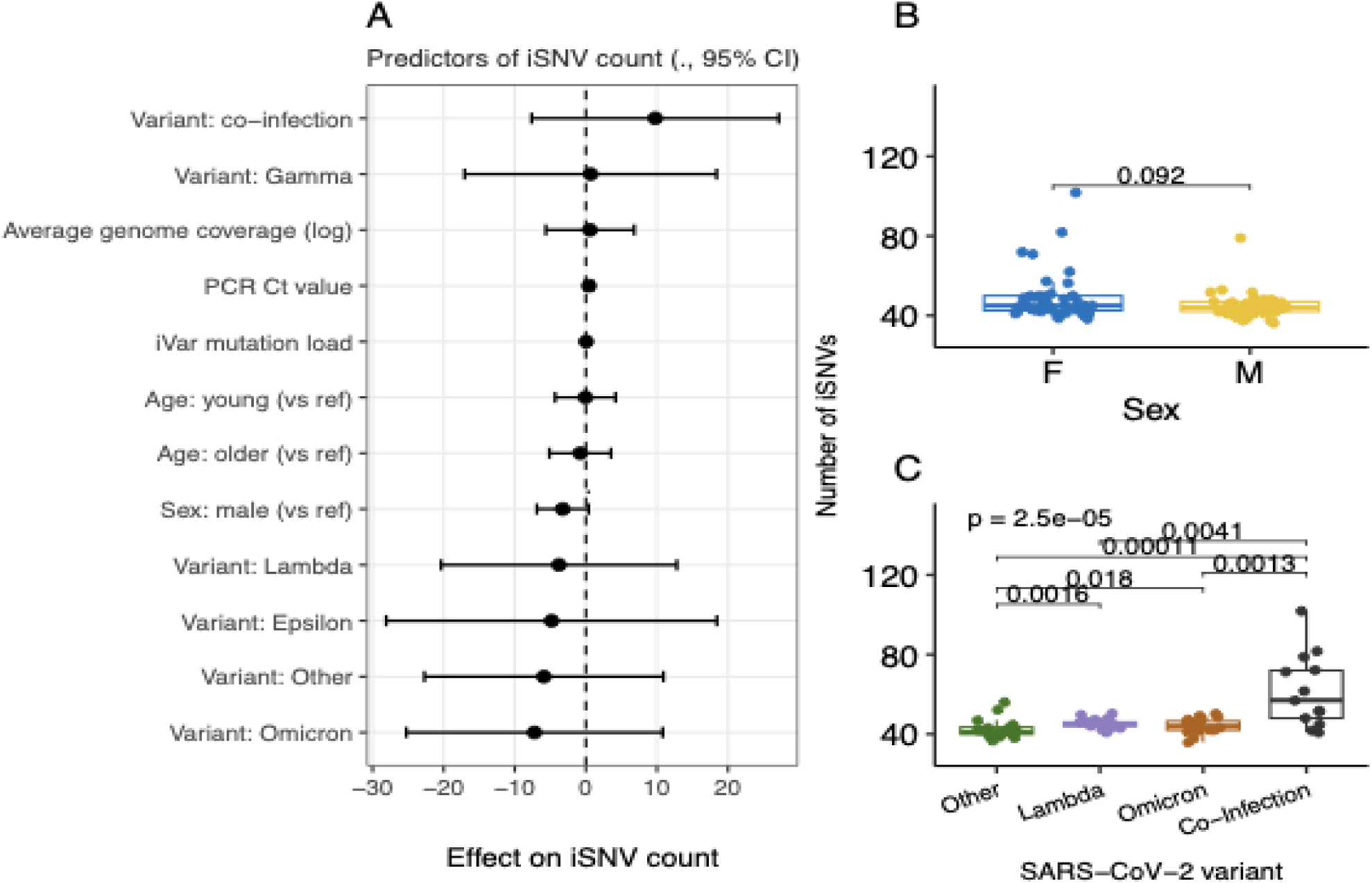
Predictors of intra-host SARS-CoV-2 single-nucleotide variants (iSNVs) in Chilean patients. (A) Multivariate linear model showing the effect size (β ± 95% CI) of demographic, molecular, and technical variables on iSNV counts. Sex and viral variant were included as categorical predictors; PCR Ct and log-transformed mean genome coverage were included as continuous covariates. (B) Comparison of iSNV counts by sex. (C) Distribution of iSNV counts among major SARS-CoV-2 variants (≥ 10 cases per group).

To probe variant iSNV differences, we compared medians among variants with ≥10 cases (Lambda, Omicron, Other, Co-Infection)(Figure 3C). A Kruskal–Wallis test indicated significant heterogeneity (p ≈ 2.5×10⁻⁵), driven by higher median iSNV counts in co-infection samples versus any single-variant group (typical medians ≈ 68 vs 42–44; Wilcoxon pairwise p ≤ 0.004 after ties-robust testing; Figure 3C). Mutational spectrum analysis of iSNVs revealed consistent transition and transversion patterns across all SARS-CoV-2 variants (Supplementary Fig. 4), with a predominance of C→T and T→C substitutions consistent with the activity of host RNA editing enzymes such as APOBEC and ADAR[24]. These enzymes are known to contribute to the genetic variability of RNA viruses by inducing deamination of cytidines (APOBEC) and adenosines (ADAR)[24]. Additionally, the G→T base pair change could be attributed to Reactive Oxygen Species (ROS)[22].

Together, these results suggest that the elevated intra-host diversity observed during co-infection events is more likely due to factors within the host, rather than sex, demographic, or technical factors.

### Selection dynamics across intra-host SARS-CoV-2 variants

We estimated nonsynonymous-to-synonymous substitution ratios (dN/dS) from iSNVs to assess the selective pressures acting on SARS-CoV-2 genomes circulating in Chile across successive epidemic waves [25]. Global dN/dS values were below 1 for all lineages, indicating that purifying selection predominated during intra-host replication[20,21,25]. Early *Other* lineages (2020) exhibited the highest overall ratio (0.63; 95% CI: 0.41–0.97), followed by *Lambda* (0.62; 0.38–1.0), *Co-infection* samples (0.58; 0.43–0.79), and *Omicron* (0.55; 0.34–0.87) (Figure 4A). This increased purifying selection over time reflects the progressive optimization of the viral genome[21]. Co-infections, most involving Omicron plus a second lineage, displayed an intermediate global burden (0.58), indicating that mixed viral populations preserve Omicron’s overall constraint while incorporating additional missense diversity contributed by the co-infecting variant[20,21].

**Figure 4.**
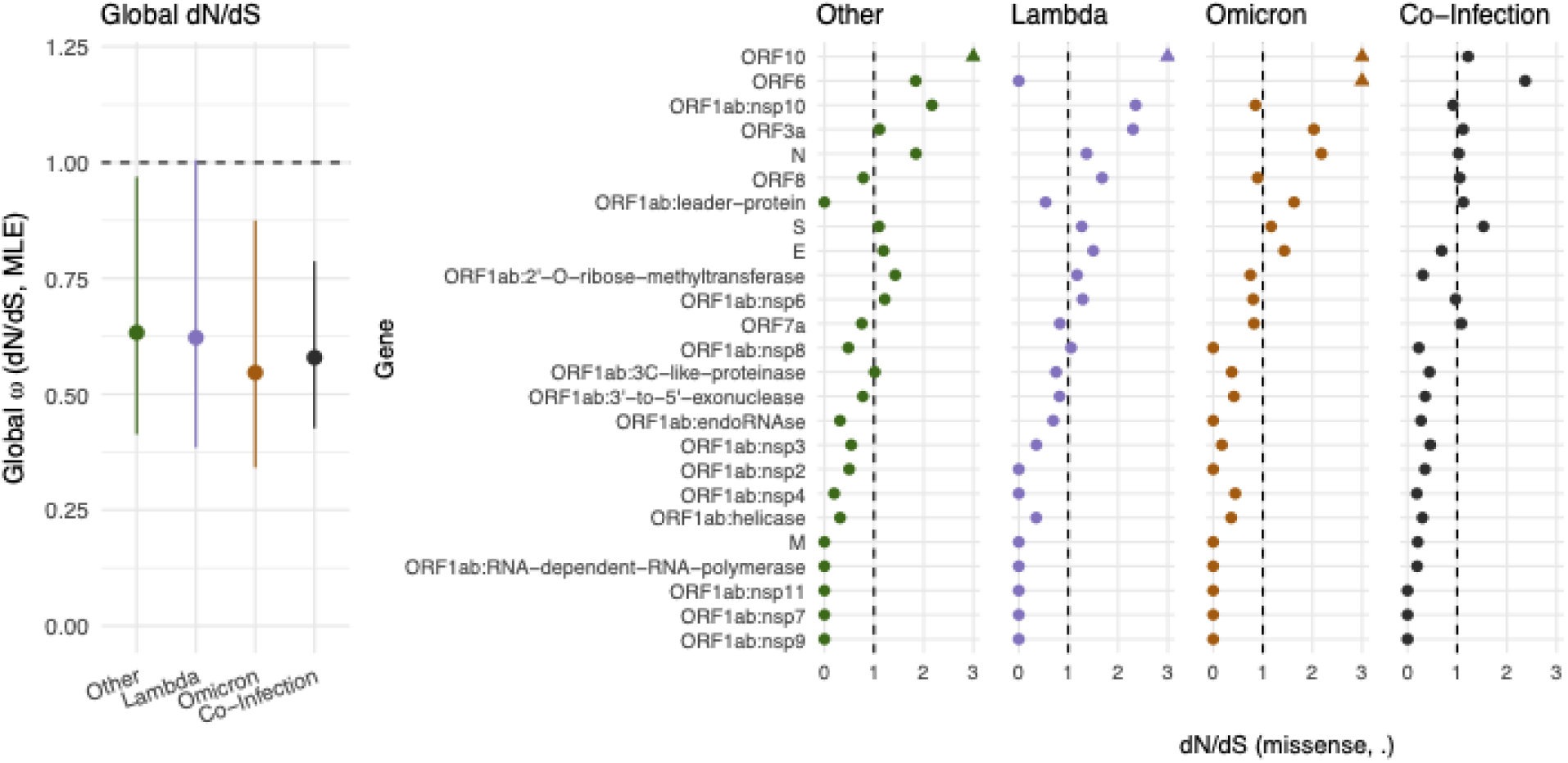
Global and Gene-level iSNV dN/dS estimates across SARS-CoV-2 variants. a) Global dN/dS (ω) estimates derived from iSNVs for each variant group: Other, Lambda, Omicron, and Co-infections. Points represent the maximum-likelihood estimate (MLE), and vertical bars show 95% confidence intervals. The dashed horizontal line marks ω = 1, the neutral expectation. b) Gene-level iSNV dN/dS values for SARS-CoV-2 genes across variant groups. Each column corresponds to a variant, and each point represents the estimated dN/dS for a specific gene. Horizontal dashed lines mark ω = 1 (neutral evolution).

We next compared gene-level dN/dS ratios derived from iSNVs across the four variant groups (Other, Lambda, Omicron, and Co-infections, Figure 4B). A Kruskal–Wallis test revealed no significant differences between groups (χ² = 0.38, df = 3, p = 0.94), and pairwise Dunn tests further confirmed the absence of variant-specific effects, with all adjusted p-values equal to 1.0 (FDR). These results indicate that the intra-host selective pressures reflected by iSNV-based dN/dS estimates are broadly conserved across SARS-CoV-2 variants, aligning with prior observations that viral evolution is shaped by localized episodic adaptation (e.g., in Spike, ORF6, ORF10, ORF8) rather than genome-wide shifts in purifying selection[3,21,25]. However, clear temporal shifts in intra-host selection targets were still apparent from the iSNV-based gene-level dN/dS estimates. In early and pre-Omicron lineages (Other and Lambda), several genes exhibited dN/dS values > 1, indicating episodic positive selection during the initial phase of viral adaptation. These included S, N, ORF6, ORF10, ORF3a, and multiple ORF1ab non-structural proteins (e.g., nsp6, nsp10, 2’-O-ribose-methyltransferase, Figure 4B). For example, the spike gene showed dN/dS values of 1.10 (Other) and 1.27 (Lambda), consistent with early diversification of receptor-binding and antigenic regions. Genes such as N (1.84 and 1.37) and ORF10 (4.27 and 5.30) also exhibited elevated ratios, reflecting selective pressures on virion assembly, immune modulation, or accessory protein function during the expansion of these lineages[3]. Following the emergence of Omicron, most genes returned to dN/dS < 1, consistent with the strong purifying selection that occurred after the fixation of major adaptive changes (Figure 4B). Nevertheless, Omicron retained modest elevation in specific loci, including N (2.18), S (1.17), ORF3a (2.03), and ORF6 (3.56), indicating continued refinement of host interaction, immune evasion, and accessory gene function during rapid global spread. Co-infection samples displayed a distinct pattern. Although dN/dS ratios were generally lower than those of early variants(Figure 4B), several genes again exhibited values > 1, including S (1.53), N (1.03), ORF3a (1.12), ORF6 (2.37), ORF10 (1.22), and multiple non-structural proteins (leader protein, nsp6, methyltransferase). This suggests transient reactivation of adaptive pressures during mixed infections, potentially driven by intra-host competition, replication interference, or opportunities for recombination that arise when multiple genomes co-replicate within the same host cell environment[3].

## Discussion

Our study presents the first ultra-deep sequencing and intra-host evolutionary analysis of SARS-CoV-2 in Chile, spanning multiple epidemic waves (2020–2022) and the transition from early variants to the dominance of Omicron in Chile. Using the ATOPlex platform and DNBSeq-G400 sequencing, we achieved an average depth of ∼60,000×, allowing high-confidence detection of low-frequency iSNVs. The resulting dataset captures the evolutionary dynamics of SARS-CoV-2 across key stages of the pandemic in Chile, complementing national genomic surveillance that previously relied primarily on consensus-level data.

Our results reveal that within-host diversity is primarily shaped by viral lineage and co-infection status, rather than host demographic or technical factors. While the number of fixed mutations (SNVs) increased over time, reflecting global lineage diversification, the overall iSNV burden remained relatively stable across the study period. Importantly, co-infection cases, which accounted for approximately 20% of our samples, exhibited markedly higher iSNV counts than single-variant infections. Most co-infections involved Omicron sublineages (BA.2* and BA.5*) coexisting with pre-Omicron variants such as Gamma and Lambda, highlighting the epidemiological overlap between circulating lineages during Chile’s 2021–2022 transition. These findings align with emerging reports that the co-circulation of divergent SARS-CoV-2 variants can lead to transient co-infections, creating conditions favorable for recombination and rapid diversification[20,26].

At the molecular level, our dN/dS analysis provides new insights into the selective dynamics of SARS-CoV-2 within hosts across successive epidemic phases in Chile. Consistent with global observations[3,20,21,25], we found that purifying selection is the dominant evolutionary force acting on the viral genome, with global dN/dS ratios remaining below 1 across all variants. Early circulating lineages, grouped as Other and Lambda, exhibited moderate dN/dS values (0.63 and 0.62, respectively), suggesting episodes of adaptive diversification during the initial viral establishment in the population. By contrast, *Omicron* showed stronger purifying selection (dN/dS = 0.55), reflecting the extensive optimization of its genome and reduced evolutionary space following multiple prior adaptive sweeps[3].

A key finding of our study is the increased dN/dS ratio in co-infection cases (0.58), accompanied by gene-specific signals of positive selection. This supports the notion that co-infection can facilitate within-host diversification and recombination, potentially accelerating the emergence of novel variants [20,26]. Our data also capture a temporal trajectory of selection spanning the Chilean epidemic, from adaptive diversification in early variants (Other and Lambda) through strong purifying constraints in *Omicron*, to renewed adaptive potential in co-infected hosts. This pattern mirrors the global evolutionary progression of SARS-CoV-2, yet it is revealed here at an intra-host scale, enabled by ultra-deep sequencing coverage that allows high-confidence detection of low-frequency alleles. The results emphasize how within-host selection dynamics mirror and sometimes precede population-level trends, suggesting that the seeds of future variant emergence may originate during co-infection events or within individuals hosting multiple lineages[3,26].

Our study offers new insights into the within-host diversity of SARS-CoV-2 in the Chilean population; however, several limitations should be acknowledged. The cohort of 96 individuals, drawn from only two regions, may not capture the full geographic and epidemiological breadth of viral diversity in Chile. Nonetheless, the ultra–high-depth sequencing applied here enabled sensitive detection of iSNVs across all samples. Future work should include larger and more geographically diverse cohorts and integrate clinical and epidemiological data to better resolve the factors shaping within-host evolution.

Together, our findings contribute to the understanding of SARS-CoV-2 evolution by showing that while purifying selection governs most within-host replication, co-infection provides transient evolutionary windows for diversification and adaptive experimentation. Monitoring such events using ultra-deep sequencing may improve our ability to detect early signals of viral innovation and anticipate the emergence of recombinant or more transmissible lineages.

## Material and Methods

### SARS-COV-2 genome sequencing using ATOPlex and DNBSeq G400

ATOPlex[27] SARS-CoV-2 sequencing is a two-step, multiplex-PCR-based Massively Parallel Sequencing (mPCR-based MPS) performed on the DNBSEQ platform[28]. We utilized ATOPlex to sequence the SARS-CoV-2 genome from nasopharyngeal (NP) swabs of 96 confirmed COVID-19 individuals. The multiplex PCR of ATOPlex utilizes short amplicons (106-199 bp) with high specificity and sensitivity, allowing for the generation of complete SARS-CoV-2 virus genomes from samples with PCR Ct values of up to 35. Library preparation was performed using the ATOPlex RNA Library Prep Set (MGI, China) according to the manufacturer’s instructions. In Briefly, 10ul total RNA of each sample was converted to the first-strand cDNA products by RNA reverse transcriptase with random hexamers (5’-NNNNNN-3’). 1st strand cDNA products were then amplified by ATOPlex SARS-CoV-2 full-length genome panel and universal barcode primer. After amplification, the second-round PCR products were quantified with the Qubit dsDNA High Sensitivity Assay kit ((Thermo Fisher Scientific, Waltham, MA, USA) to confirm the required concentration of ≥4 ng/µL Short amplicons libraries from each sample were pooled at equimolar levels and subjected to single-stranded circular DNA library preparation with the MGIEasy Dual Barcode Circularization kit (MGI) to obtain circularized DNA molecules. These molecules were subsequently digested to form circularized single-strand DNA (ssCirDNA) and then subjected to rolling circle amplification to generate DNA nanoball (DNB) libraries. The DNBs were added to a silicon slide that contains a grid-like pattern of binding sites, which enables the DNBs to self-assemble into a dense grid of spots for sequencing. The DNB libraries were sequenced in-house on a DNBSEQ-G400 instrument with paired-end reads of 150 base pairs. Sequencing was performed in-house at the University of O’Higgins.

### SARS-COV-2 sequence analysis

We used the CECRET pipeline(version 0fdad331), implemented in Nextflow[31], and it integrates a suite of bioinformatics tools for comprehensive analysis of SARS-CoV-2 sequencing data[30]. We executed the pipeline using containers and the following command: *nextflow run UPHL-BioNGS/Cecret -profile singularity --cleaner fastp --relatedness true --samtools_ampliconstats false --samtools_plot_ampliconstats false.* In brief, this command and its options initiate the CECRET pipeline, which cleans the reads using fastp[31], followed by the generation of quality control metrics with FastQC. The reads were then aligned to the reference genome Wuhan-Hu-1 (RefSeq ID NC_045512) using BWA-MEM[35], and processed with SAMtools[36] for sorting, indexing, and various QC metrics. Variant calling was performed using BCFtools [30], and iVar [31] was employed for variant calling, creating consensus FASTAs, and primer trimming. MAFFT was employed for multiple sequence alignment, and phylogenetic tree generation was performed using IQ-TREE2[37]. Nextclade[38] and Pangolin[39] were used for SARS-CoV-2 clade classification. Freyja[40] was used to recover relative lineage abundances from SARS-CoV-2 samples, which were aligned to the Wuhan-Hu-1 reference using the BAM format[36]. Finally, MultiQC[41] was run to summarize all previous results. The CECRET pipeline execution took 320 CPU hours on the kutral cluster of the HPC-UOH infrastructure at Universidad de O’ Higgins.

### iSNVs analysis

To identify high-confidence intra-host single-nucleotide variants (iSNVs) from ultra-deep sequencing, we applied stringent filtering criteria requiring an alternative allele depth of at least 100 reads, a Phred quality score >30, and an allele frequency between 5% and 95% to remove low-frequency artefacts and fixed consensus variants. Positions overlapping sample-specific consensus SNVs were excluded. To ensure sufficient viral template input, only samples with Ct ≤28 were retained for downstream analyses. For mutation spectrum analysis, substitution types were extracted from high-confidence iSNVs and summarized across samples to quantify the frequencies of transitions and transversions. To evaluate demographic, molecular, and technical predictors of intra-host diversity, we modeled the number of iSNVs per sample using a multivariate linear regression model in R. Model fit was evaluated using R², adjusted R², and F-statistics, and effect sizes were plotted with 95% confidence intervals to visualize parameter uncertainty.

Co-infections were identified by integrating iVar iSNV profiles with lineage deconvolution using Freyja[40], which was run on BAM alignments to estimate relative lineage abundances. Samples exhibiting at least two distinct viral lineages, each contributing ≥3% abundance, were classified as putative co-infections. To compare iSNV burdens among major viral variants represented by at least 10 samples (Other, Lambda, Omicron, and Co-infection), we used the Kruskal–Wallis test followed by pairwise Wilcoxon rank-sum tests with ties-robust p-value estimation. Median iSNV counts were reported for each variant group. Intra-host selective pressures were quantified by estimating the nonsynonymous-to-synonymous substitution rate ratio (dN/dS) using the dndscv framework [42], applied independently to each variant group. Global dN/dS values and gene-level estimates were obtained from maximum-likelihood models that account for mutational signatures and codon structure. Differences in gene-level selection across variants were assessed using Kruskal–Wallis tests followed by Dunn post-hoc comparisons with FDR correction. All statistical analyses and figures were carried out using R.

## Data and code availability

The dataset and code used in the main analyses, along with the figures generated for this work, are available on GitHub (https://github.com/digenoma-lab/covid_genomics). Raw sequencing reads were submitted to SRA under the bioproject XXXX.

## Author Contributions

Conceptualization: ML, ADG; Methodology: AG, CM, ML, ADG; Investigation: FK, BB, SP, PJ, CM, ML, ADG; Resources: AG, CM, ML, ADG; Visualization: FK, BB, SP, PJ, ADG; Funding Acquisition: ML, ADG; Project Administration: ADG; Supervision: CM, ML, ADG; Writing – Original Draft: CM, FK, BB, SP, PJ, ADG; Writing – Review & Editing: FK, CM, ML, ADG. All authors have read and agreed to the published version of the manuscript.

## Anknowledge

We thank Professor Ricardo Andres Soto from Universidad de Chile for the critical reading and helpful comments on our manuscript.

## References

1. [cited 2 Dec 2025]. Available: https://data.who.int/dashboards/covid19/case

2. Telenti A, Arvin A, Corey L, Corti D, Diamond MS, García-Sastre A, et al. After the pandemic: perspectives on the future trajectory of COVID-19. Nature. 2021;596: 495–504.

3. Markov PV, Ghafari M, Beer M, Lythgoe K, Simmonds P, Stilianakis NI, et al. The evolution of SARS-CoV-2. Nat Rev Microbiol. 2023;21: 361–379.

4. Gao L, Zheng C, Shi Q, Xiao K, Wang L, Liu Z, et al. Evolving trend change during the COVID-19 pandemic. Front Public Health. 2022;10: 957265.

5. Islam MR. The SARS-CoV-2 Omicron (B.1.1.529) variant and the re-emergence of COVID-19 in Europe: An alarm for Bangladesh. Health Sci Rep. 2022;5: e545.

6. McCallum M, Czudnochowski N, Rosen LE, Zepeda SK, Bowen JE, Walls AC, et al. Structural basis of SARS-CoV-2 Omicron immune evasion and receptor engagement. Science. 2022;375: 864–868.

7. Arora P, Zhang L, Rocha C, Sidarovich A, Kempf A, Schulz S, et al. Comparable neutralisation evasion of SARS-CoV-2 omicron subvariants BA.1, BA.2, and BA.3. Lancet Infect Dis. 2022;22: 766–767.

8. Pulliam JRC, van Schalkwyk C, Govender N, von Gottberg A, Cohen C, Groome MJ, et al. Increased risk of SARS-CoV-2 reinfection associated with emergence of Omicron in South Africa. Science. 2022;376: eabn4947.

9. Gerli AG, Centanni S, Soriano JB, Ancochea J. Forecasting COVID-19 infection trends in the EU-27 countries, the UK and Switzerland due to SARS-CoV-2 Variant of Concern Omicron. bioRxiv. 2021. doi:10.1101/2021.12.16.21267785

10. Jara A, Undurraga EA, Zubizarreta JR, González C, Acevedo J, Pizarro A, et al. Effectiveness of CoronaVac in children 3-5 years of age during the SARS-CoV-2 Omicron outbreak in Chile. Nat Med. 2022;28: 1377–1380.

11. Jara A, Cuadrado C, Undurraga EA, García C, Nájera M, Bertoglia MP, et al. Effectiveness of the second COVID-19 booster against Omicron: a large-scale cohort study in Chile. Nat Commun. 2023;14: 6836.

12. Nogareda F, Regan AK, Couto P, Fowlkes AL, Gharpure R, Loayza S, et al. Effectiveness of COVID-19 vaccines against hospitalisation in Latin America during three pandemic waves, 2021-2022: a test-negative case-control design. Lancet Reg Health Am. 2023;27: 100626.

13. Méndez C, Peñaloza HF, Schultz BM, Piña-Iturbe A, Ríos M, Moreno-Tapia D, et al. Humoral and cellular response induced by a second booster of an inactivated SARS-CoV-2 vaccine in adults. EBioMedicine. 2023;91: 104563.

14. Mena GE, Martinez PP, Mahmud AS, Marquet PA, Buckee CO, Santillana M. Socioeconomic status determines COVID-19 incidence and related mortality in Santiago, Chile. Science. 2021;372. doi:10.1126/science.abg5298

15. Hodcroft EB, Zuber M, Nadeau S, Vaughan TG, Crawford KHD, Althaus CL, et al. Spread of a SARS-CoV-2 variant through Europe in the summer of 2020. Nature. 2021;595: 707–712.

16. GISAID - gisaid.org. [cited 1 Dec 2025]. Available: https://gisaid.org/

17. Berno G, Fabeni L, Matusali G, Gruber CEM, Rueca M, Giombini E, et al. SARS-CoV-2 Variants Identification: Overview of Molecular Existing Methods. Pathogens. 2022;11. doi:10.3390/pathogens11091058

18. Harper H, Burridge A, Winfield M, Finn A, Davidson A, Matthews D, et al. Detecting SARS-CoV-2 variants with SNP genotyping. PLoS One. 2021;16: e0243185.

19. Wang Y, Wang D, Zhang L, Sun W, Zhang Z, Chen W, et al. Intra-host variation and evolutionary dynamics of SARS-CoV-2 populations in COVID-19 patients. Genome Med. 2021;13: 30.

20. Li J, Du P, Yang L, Zhang J, Song C, Chen D, et al. Two-step fitness selection for intra-host variations in SARS-CoV-2. Cell Rep. 2022;38: 110205.

21. Lythgoe KA, Hall M, Ferretti L, de Cesare M, MacIntyre-Cockett G, Trebes A, et al. SARS-CoV-2 within-host diversity and transmission. Science. 2021;372. doi:10.1126/science.abg0821

22. Gu H, Quadeer AA, Krishnan P, Ng DYM, Chang LDJ, Liu GYZ, et al. Within-host genetic diversity of SARS-CoV-2 lineages in unvaccinated and vaccinated individuals. Nat Commun. 2023;14: 1793.

23. Ren X, Jin Q. Safe sequencing depth to estimate the intra-host heterogeneity of viruses. Brief Funct Genomics. 2016;15: 275–277.

24. Di Giorgio S, Martignano F, Torcia MG, Mattiuz G, Conticello SG. Evidence for host-dependent RNA editing in the transcriptome of SARS-CoV-2. Sci Adv. 2020;6: eabb5813.

25. Tonkin-Hill G, Martincorena I, Amato R, Lawson ARJ, Gerstung M, Johnston I, et al. Patterns of within-host genetic diversity in SARS-CoV-2. Elife. 2021;10. doi:10.7554/eLife.66857

26. Pipek OA, Medgyes-Horváth A, Stéger J, Papp K, Visontai D, Koopmans M, et al. Systematic detection of co-infection and intra-host recombination in more than 2 million global SARS-CoV-2 samples. Nat Commun. 2024;15: 517.

27. Ni G, Lu J, Maulani N, Tian W, Yang L, Harliwong I, et al. Novel multiplexed amplicon-based sequencing to quantify SARS-CoV-2 RNA from wastewater. Environ Sci Technol Lett. 2021;8: 683–690.

28. Jeon SA, Park JL, Park S-J, Kim JH, Goh S-H, Han J-Y, et al. Comparison between MGI and Illumina sequencing platforms for whole genome sequencing. Genes Genomics. 2021;43: 713–724.

29. Ahmed W, Bivins A, Metcalfe S, Smith WJM, Ziels R, Korajkic A, et al. RT-qPCR and ATOPlex sequencing for the sensitive detection of SARS-CoV-2 RNA for wastewater surveillance. Water Res. 2022;220: 118621.

30. Cecret: Reference-based consensus creation. Github; Available: https://github.com/UPHL-BioNGS/Cecret

31. Di Tommaso P, Chatzou M, Floden EW, Barja PP, Palumbo E, Notredame C. Nextflow enables reproducible computational workflows. Nat Biotechnol. 2017;35: 316–319.

32. Rao SN, Manissero D, Steele VR, Pareja J. A systematic review of the clinical utility of cycle threshold values in the context of COVID-19. Infect Dis Ther. 2020;9: 573–586.

33. Mushegian AA, Long SW, Olsen RJ, Christensen PA, Subedi S, Chung M, et al. Within-host genetic diversity of SARS-CoV-2 in the context of large-scale hospital-associated genomic surveillance. medRxiv. 2022. doi:10.1101/2022.08.17.22278898

34. Takahashi T, Ellingson MK, Wong P, Israelow B, Lucas C, Klein J, et al. Sex differences in immune responses that underlie COVID-19 disease outcomes. Nature. 2020;588: 315–320.

35. Li H. Aligning sequence reads, clone sequences and assembly contigs with BWA-MEM. arXiv [q-bio.GN]. 2013. Available: http://arxiv.org/abs/1303.3997

36. Danecek P, Bonfield JK, Liddle J, Marshall J, Ohan V, Pollard MO, et al. Twelve years of SAMtools and BCFtools. Gigascience. 2021;10. doi:10.1093/gigascience/giab008

37. Minh BQ, Schmidt HA, Chernomor O, Schrempf D, Woodhams MD, von Haeseler A, et al. IQ-TREE 2: New Models and Efficient Methods for Phylogenetic Inference in the Genomic Era. Mol Biol Evol. 2020;37: 1530–1534.

38. Aksamentov I, Roemer C, Hodcroft E, Neher R. Nextclade: clade assignment, mutation calling and quality control for viral genomes. J Open Source Softw. 2021;6: 3773.

39. O’Toole Á, Scher E, Underwood A, Jackson B, Hill V, McCrone JT, et al. Assignment of epidemiological lineages in an emerging pandemic using the pangolin tool. Virus Evol. 2021;7: veab064.

40. Karthikeyan S, Levy JI, De Hoff P, Humphrey G, Birmingham A, Jepsen K, et al. Wastewater sequencing reveals early cryptic SARS-CoV-2 variant transmission. Nature. 2022;609: 101–108.

41. Ewels P, Magnusson M, Lundin S, Käller M. MultiQC: summarize analysis results for multiple tools and samples in a single report. Bioinformatics. 2016;32: 3047–3048.

42. Martincorena I, Raine KM, Gerstung M, Dawson KJ, Haase K, Van Loo P, et al. Universal Patterns of Selection in Cancer and Somatic Tissues. Cell. 2017;171: 1029–1041.e21.

