## Supplementary material 1 for "Ultra-deep sequencing reveals intra-host diversity and co-infection-driven evolution of SARS-CoV-2"

**Supplementary Table 1. Summary of SARS-CoV-2 co-infections detected in Chilean samples**. Samples were grouped by sequencing file, and variants contributing more than 3% of the total signal were retained for further analysis. The number of unique variants and their composition are shown for each sample. All listed samples met the co-infection criterion (≥ 2 distinct variants with > 3% abundance). The most common co-infecting lineages included Omicron (BA.2* or BA.5*) with Other, Gamma, or Lambda variants, consistent with overlapping epidemic waves between 2021 and 2022.

| Sample ID | Number of Variants | Detected Variants (> 3%) |
|---|---|---|
| 68 | 5 | Other, Gamma, BA.2* [Omicron (BA.2.X)], Omicron, Lambda |
| 88 | 5 | Omicron, BA.2* [Omicron (BA.2.X)], BA.5* [Omicron (BA.5.X)], Other, Gamma |
| 56 | 4 | BA.2* [Omicron (BA.2.X)], Other, Gamma, Omicron |
| 59 | 4 | BA.2* [Omicron (BA.2.X)], Lambda, Omicron, Other |
| 03 | 3 | Other, Omicron, BA.2* [Omicron (BA.2.X)] |
| 50 | 3 | BA.2* [Omicron (BA.2.X)], Other, Lambda |
| 53 | 3 | Other, Gamma, Lambda |
| 61 | 3 | Other, Omicron, Gamma |
| 73 | 3 | Other, Omicron, Lambda |
| 76 | 3 | Omicron, Other, Gamma |
| 77 | 3 | Other, Gamma, BA.2* [Omicron (BA.2.X)] |
| 79 | 3 | BA.2* [Omicron (BA.2.X)], Other, Omicron |
| 17 | 2 | Other, Gamma |
| 44 | 2 | Gamma, Other |
| 48 | 2 | Gamma, Other |
| 49 | 2 | BA.2* [Omicron (BA.2.X)], Other |
| 58 | 2 | BA.2* [Omicron (BA.2.X)], Other |
| 70 | 2 | Omicron, Lambda |
| 87 | 2 | BA.2* [Omicron (BA.2.X)], Lambda |

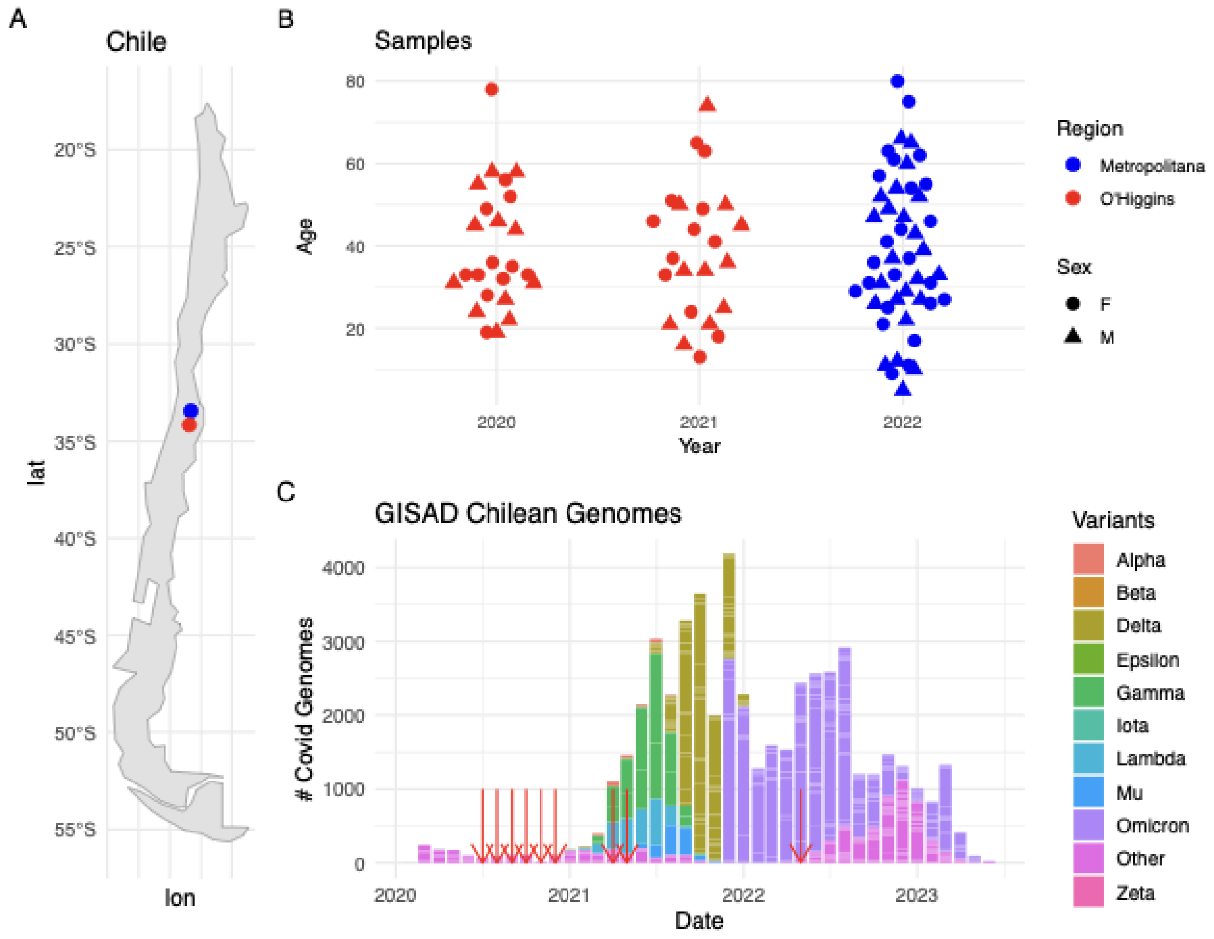


**Figure S1**: **Overview of COVID-19 Sample Sequencing**. A) Geographic Origin of Samples: The samples were collected from two regions in Chile: O'Higgins and the Metropolitan capital, Santiago. B) Demographic and Temporal Distribution of samples. C) Integration with GISAD Data: The red arrows mark the collection dates of our sequenced samples against the backdrop of Chilean genomes submitted to GISAID from 2020 to 2023 (45466 genomes at 22-09-23). This juxtaposition highlights the contribution of our data to Chilean surveillance efforts.

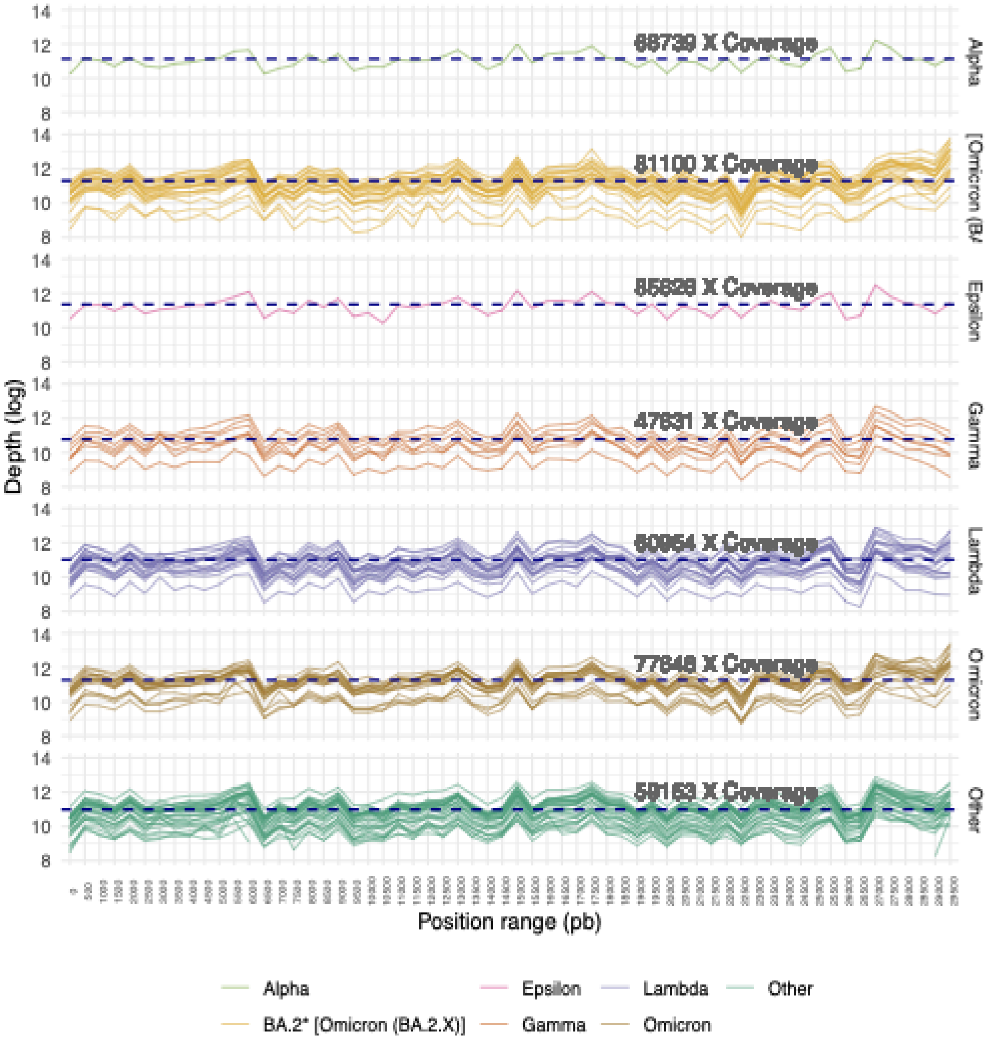


**Figure S2.** Sequencing depth of samples by the position of the genome. Samples are separated by variant (colors) and patient (multiple curves). Depth value is represented as media in windows of 0.5K pb.

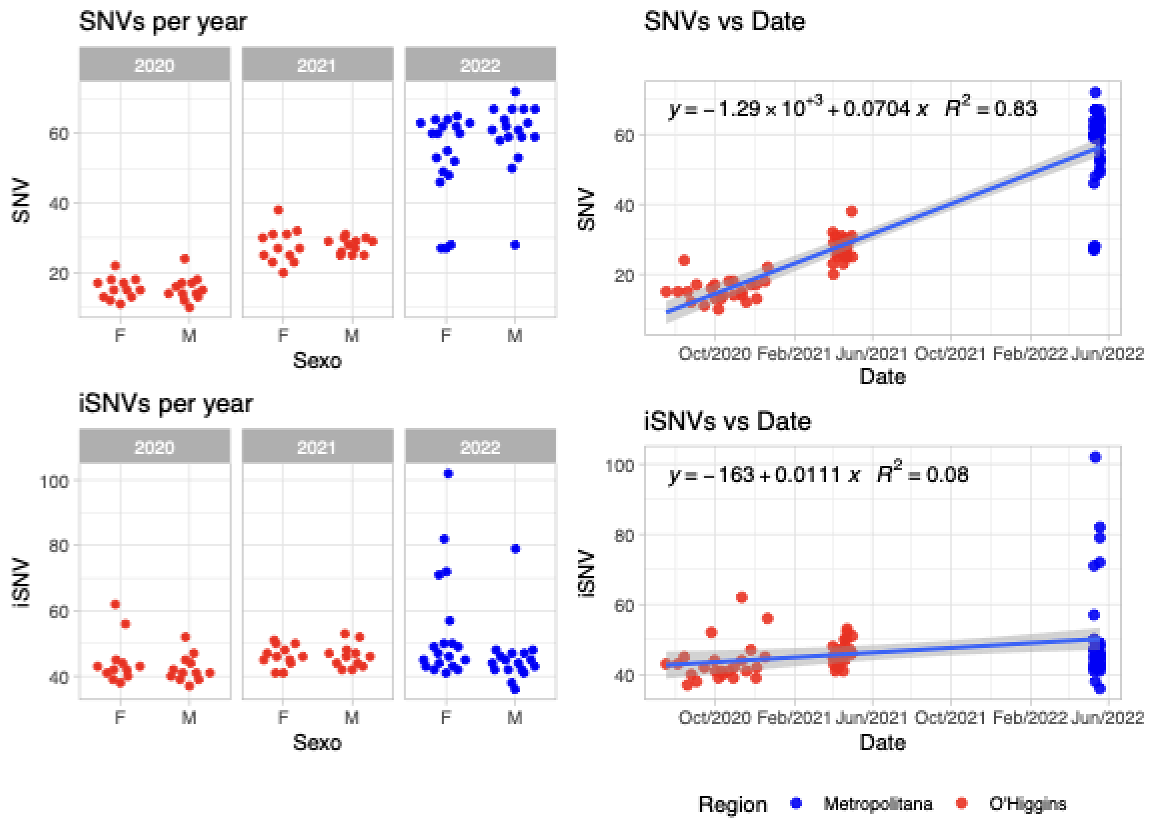


**Figure S3. Single Nucleotide Variants (SNVs) and intra-Host Single Nucleotide Variants (iSNVs) comparison by Year, Sex, and Region of Collection**. The top panels show SNVs per year (2020–2022) categorized by sex (F: Female, M: Male) with data from two regions: Metropolitana (blue) and O'Higgins (red), alongside a regression analysis revealing a strong positive correlation between SNVs and collection date ($R^2 = 0.88$). The bottom panels present iSNVs per year under the same categories, with a weaker correlation between iSNVs and collection date ($R^2 = 0.08$).

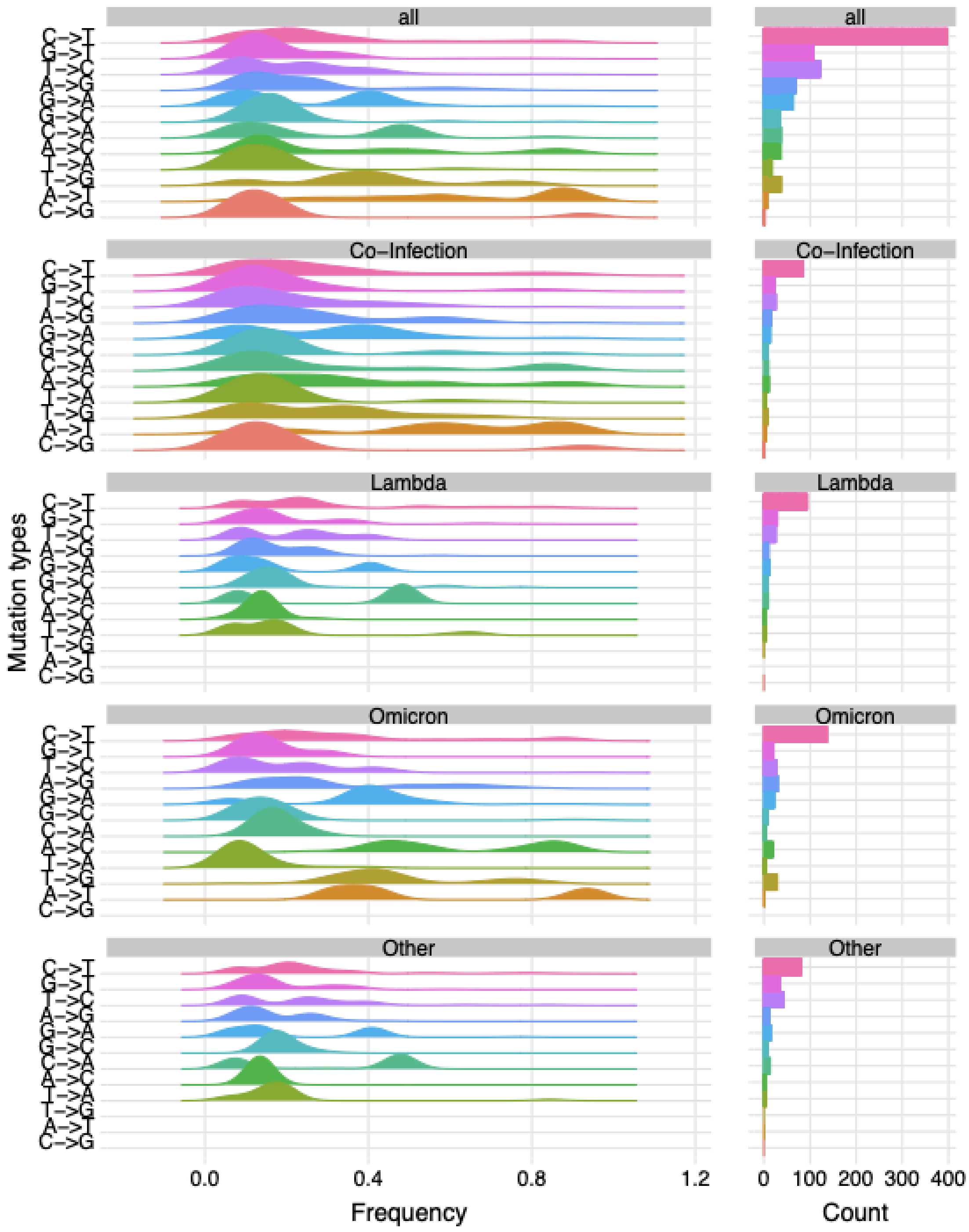


**Figure S4: Base pair change frequency for iSNVs.** Relative frequencies of transition and transversion types (A→G, C→T, G→A, etc.) identified as intra-host single-nucleotide variants (iSNVs) are shown for each major variant group (Lambda, Omicron, Co-infection, and Other) and across the full dataset.
